# Characterisation of an evolutionarily conserved RNaseH-like domain in a domesticated transposase-derived protein

**DOI:** 10.64898/2026.09.03.748810

**Authors:** Aditi Saha, Sharmistha Majumdar

## Abstract

The RNase-H fold is an ancient protein fold found in diverse nucleases that is characterised by a conserved structural core (five β strands and *a* helices), canonical DDE/D catalytic residues and catalytic mechanism. This study focuses on the evolutionary conservation of the RNaseH-like domain in the transposase-derived THAP9 family among different taxonomic groups. Phylogenetic analysis demonstrates that the structural framework required for catalysis has been maintained over the course of evolution, while other peripheral regions have undergone lineage-specific diversification. This is exemplified by a Lys residue, which is conserved across mammals, can be structurally superposed on a catalytic Asp residue found in active transposases, but is not important for DNA integration. Further, the importance of conserved motifs (YREK and YKEFR motifs) near the THAP9 catalytic residues are confirmed biochemically by performing site directed mutagenesis followed by functional assays which demonstrate that substitution mutants have decreased ability to integrate DNA. Overall, these results emphasize that DNA integration activity is controlled by interactions with several structural elements rather than by individual catalytic residues.

## Introduction

The RNase H-like domain is an evolutionarily related yet divergent structural domain that occurs across a diverse superfamily of proteins with distinct functional roles. The RNase-H fold is considered to be one of the most ancient protein folds which originated from viruses but has been adapted by diverse proteins in all kingdoms of life. RNase H-like domain was first discovered in Ribonucleases H to catalyse the cleavage of RNA from RNA-DNA hybrid in retroviruses, and hence the name RNase H (Hausen and Stein 1970). Similar domains have later been found in DNA transposases, Holliday junction resolvases, Piwi-Argonaute nucleases, in several endonucleases, Prp8 (which is the master regulator of spliceosome machinery), etc (Majorek et al. 2014).

The catalytic core of the RNase H-like domain is made of β sheets, where five β strands (β1–β2– β3–β4–β5) which occur sequentially in the primary sequence, adopt a β3–β2–β1–β4–β5 arrangement within the three-dimensional structure, giving rise to the characteristic “32145” topology of the RNase H-like fold in which β2 is antiparallel to others. These β sheets are surrounded by α helices, whose number can vary in individual domains (Fig 1). The C-terminal α helix is often preceded by an Insertion domain (Majorek et al. 2014).

**Fig 1:**
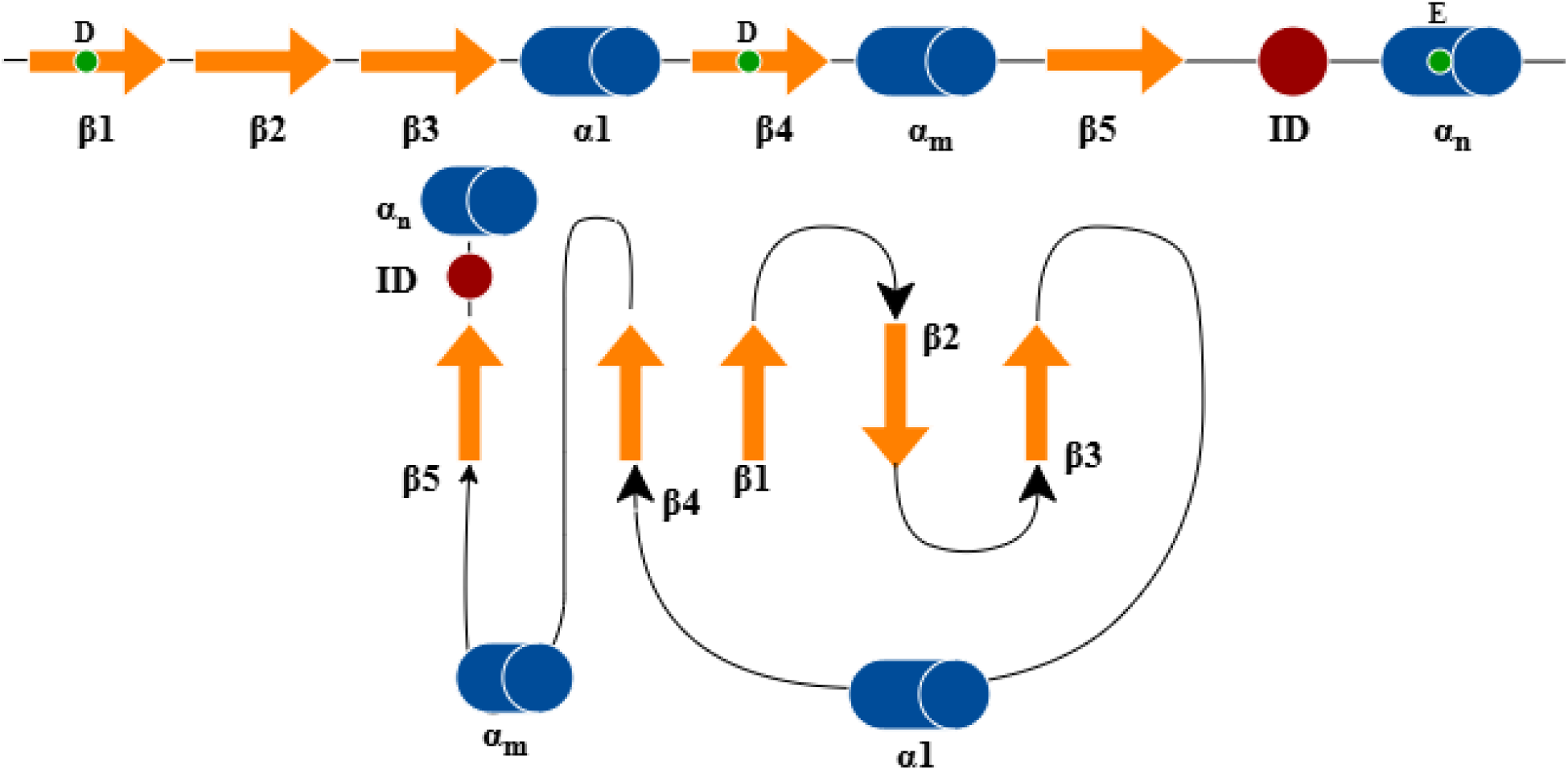
Domain architecture of RNase H-like domain for DNA transposases where the orange arrow indicate β sheet, blue cylinder indicates α helix, m and n refers to the no. of α-helices which can vary in different proteins, ID represents Insertion domain and green circles represent the DDE motif.

Insertion domains (ID) contain insertion sequences which are a characteristic feature of the catalytic domain of some RNase H-like domains in DNA transposases, recombinases, retroviral integrases (Majorek et al. 2014). For example, in Tn5, the ID is predominantly β-stranded and contains a conserved W298 that interacts with the flipped-out DNA base and facilitates hairpin formation (Davies et al. 2000; Steiniger-White et al. 2004). The absence of an insertion in transposases such as MuA, Mos1, and HIV-1 integrase is consistent with differences in second-strand processing mechanisms during transposition (Alison B. Hickman et al. 2010). In contrast, the Hermes transposase contains a larger, entirely α-helical ID that contributes both to hairpin formation and transposase oligomerization, with a conserved W319 positioned near the catalytic DDE residues (Alison B Hickman et al. 2005). RAG1 is similarly predicted to contain a large α-helical insertion between the second acidic and third acidic residues of its DDE motif, including a conserved W893, although its structural organization and precise role remain unresolved (Zhou et al. 2004; Grundy et al. 2007). These differences suggest that IDs have evolved distinct structural architectures and functions according to the mechanistic requirements of individual transposition or recombination systems. In addition, a short insertion in Mos1 between β1 and β2 (different from conventional insertion between β5 and α helix) becomes ordered upon DNA binding and contributes to transposase dimerization and protein–DNA interactions, further illustrating how insertions within the conserved RNase H-like scaffold can acquire lineage-specific structural and functional roles (Richardson et al. 2009).

RNase H-like catalytic domains contain DDE/D motifs often referred to as the “catalytic triad” which contain important acidic residues (Asp, Glu) that play a role in catalysis i.e. nucleic acid cleavage.The RNase H-like fold provides a structural scaffold that positions the conserved DDE/D catalytic residues responsible for its activity. The positions of the DD of DDE/D motif always fall on folds of β1 and β4 and the third D/E on or close to the α-helix. The presence of a conserved K/R residue at a characteristic distance downstream of the catalytic E residue provides additional support for the assignment of the RNase H-like fold. Interestingly, some IS transposases possess N or H as the third catalytic residue instead of E/D (Alison Burgess Hickman et al. 2010).

Although the detailed catalytic mechanisms differ among transposase superfamilies, they share a common mechanistic framework - the transposases having a common structural fold bring the DDE/D residues to close proximity for catalysis, coordinated by two divalent metal ions (typically Mg²⁺ or Mn²⁺). The reaction starts when the transposase enzyme binds to the TIRs of the transposon DNA followed by the hydrolysis of phosphodiester bonds leading to the formation of free 3′-OH groups in the DNA. These hydroxyl groups are protected by the transposase until they are inserted into the new target DNA via a transesterification reaction, also catalyzed by the transposase. Though the formation and architecture of the strand transfer complex varies among different transposase families, the fundamental principles of catalysis are similar (Nesmelova and Hackett 2010).

Additionally, different superfamilies of transposases (Yuan and Wessler 2011) also contain different signature strings. For example, some dsDNA transposases (namely IS231, ISH8, IS4Sa, IS4, ISPepr1, IS10 and IS50) also contain a Y-(2)-R-(3)-E-(6)-K motif, or the “YREK’ motif, which was originally discovered in the α4 helix of IS4, that belongs to the prokaryotic IS family of transposases (De Palmenaer et al. 2008). The E of the YREK motif coincides with that of the DD<u>E</u> motif of the catalytic triad. The side chains of Y and R are important for hairpin resolution while K is important for synaptic complex formation (Naumann and Reznikoff 2002). The presence of two Trp (W) residues before and after R also helps in ‘push and pull’ configuration of Tn5 (member of IS4 family); here the W323 residue after R322 drives the thymidine base of the non-transferred stand to flip out of the DNA helix and stacks against the indole ring of W298. On the other hand, Tn10 transposase (member of IS4 family) has a Met after R of the YREK motif that has a similar role in base extrusion and flipping (J. Bischerour and Chalmers 2007; J. Bischerour and Chalmers 2009).

The catalytic mechanisms of eukaryotic transposases, like *piggyBac,* proceed through hairpin loop formation, but the mechanism is neither via the ‘push and pull’ mechanism of Tn5 nor by the presence of conserved YREK residues. Other eukaryotic transposase-derived proteins like Rag do not have the YREK motif, but have a YK<u>E</u>RFK motif surrounding the E residue of the DDE catalytic triad, which has functional similarities to the prokaryotic YREK motif. Further, the presence of a W residue near the YKEFRK motif suggests that it may be involved in hairpin formation, though the exact role is unclear (Lu et al. 2008).

Human THAP9 (hTHAP9) is a homologue of the active *Drosophila* P-element transposase (DmTNP), whose exact function is unknown. Although it is assumed that THAP9 is possibly domesticated since it does not possess canonical TIRs and TSDs, it retains its catalytic activity and can mobilise the terminal inverted repeats (TIRs) of *Drosophila* P-element.

The cryo EM structure of DmTNP shows that its RNase H-like domain has a canonical RNase H-like fold with a conserved DDE motif (D230, D303, E531) coordinated by Mg^2+^ ions. Moreover, an α-helical GTP binding insertion domain is also present between β5 and α4. GTP is not necessary as an energy driving force but serves as a cofactor during the strand transfer process (Ghanim et al. 2019). The RNase H-like domain of hTHAP9 also has the conventional RNaseH-like fold and retains the ability to cut the *Drosophila* TIRs (Sharma et al. 2021). The structural superposition of the RNase H-like domains of DmTNP and hTHAP9 illustrates that the DDE (D230, D303, E531) of DmTNP aligns with K282, D374, E613 of hTHAP9. Thus, hTHAP9 appears to have a D and E corresponding to the second and the third catalytic residues of DmTNP. It is worth exploring whether the K of hTHAP9 that overlaps with DmTNP’s 1st catalytic D, plays any role in mobilisation of *Drosophila* P-element TIRs.

Further, hTHAP9 has both the prokaryotic YREK as well as YKEFR (without the last K) residues on either end of the RNase-H-like catalytic domain (boundaries predicted by AlphaFold) with the YKEFR motif carrying the E that overlaps with the DmTNP catalytic triad (Fig 3).

**Fig 2:**
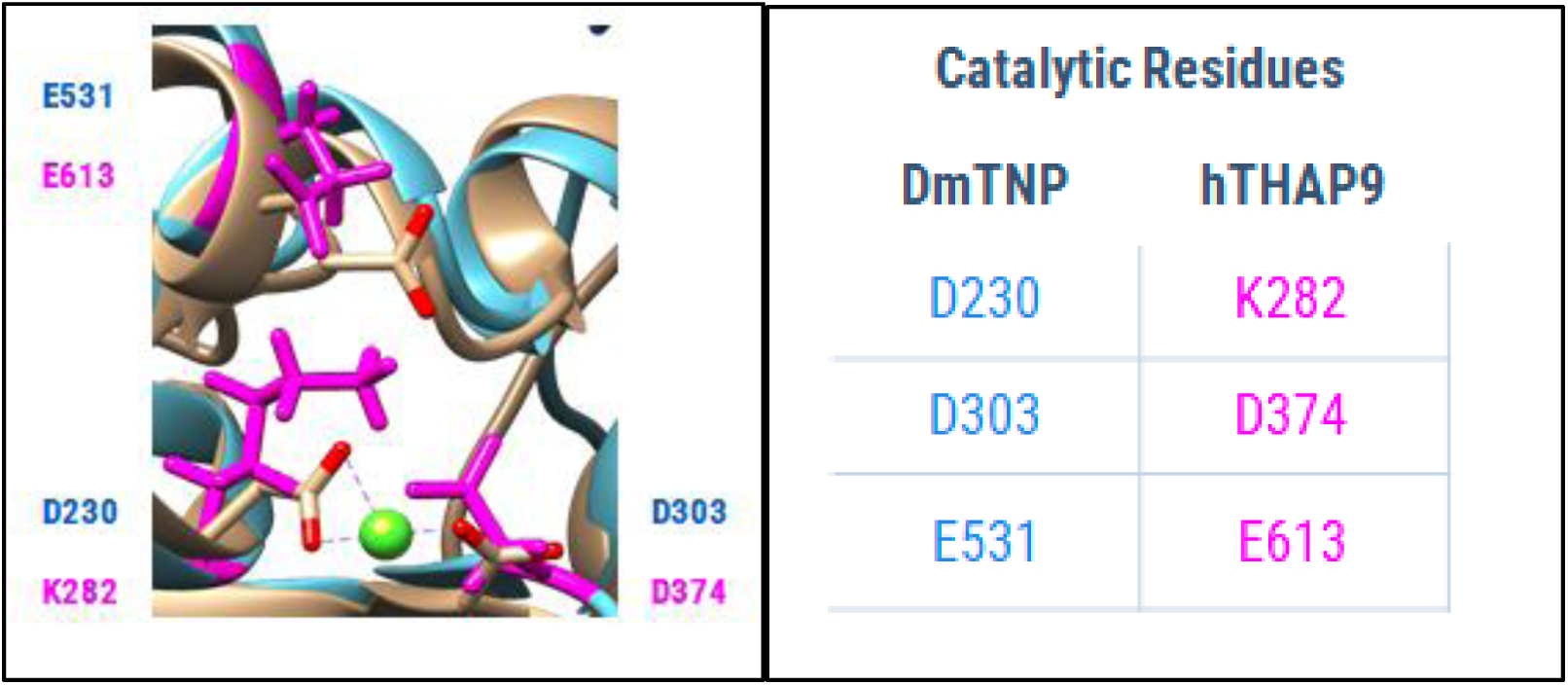
Superposition of DDE catalytic triad of DmTNP with KDE of hTHAP9 (Rashmi 2022).

**Fig 3:**
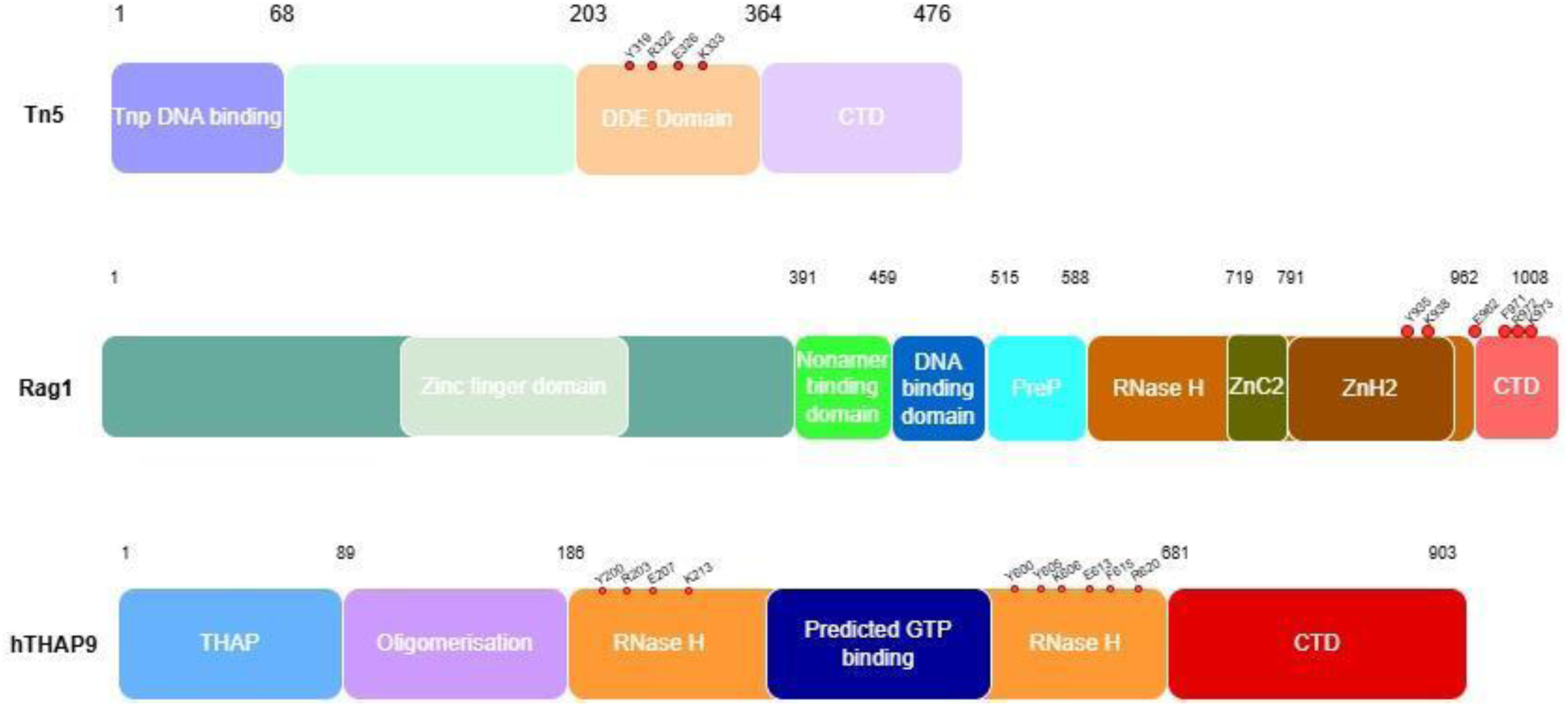
Domain architectures of transposons having only YREK domain (Tn5), only YKEFRK (Rag1), and both the YREK and YKEFR domains (hTHAP9)

Here we carry out a detailed evolutionary analysis of the RNase H-like domain of hTHAP9 to determine the extent of its conservation and divergence across different taxonomic groups, identify conserved residues (like the DDE, YREK, YKEFRK motifs) and their class-specificity, and assess how the catalytic core and its associated insertions have evolved. This analysis will provide a broader evolutionary framework for understanding the structural organization and functional diversification of the RNase H-like domain in THAP9. Further, biochemical investigations of uncharacterised conserved motifs (namely KDE, YREK, YKEFRK) in the RNaseH-like domain of hTHAP9 were carried out to understand the role of these residues in catalysis i.e. DNA cleavage and subsequent integration.

## Material and Methods

### Data collection

The amino acid sequences of RNase H-like domains (residues 181-681 in hTHAP9) present in individual homologs of hTHAP9 were taken from the InterPro database, which had only one reviewed entry (Q9H5L6, THAP9 *Homo sapiens*)(Blum et al. 2025). To obtain homologous sequences of this RNAse H-like domain, the search was performed using phmmer (Rajković et al. 2026). The 11466 retrieved sequences were obtained, keeping the database as UniProt (accessed in April 2026), and e-values and Hit-Scores as 1. Multiple sequence alignment with these sequences served as input to carry out hmmbuild (Rajković et al. 2026). The resultant hmmprofile was then used as a query to identify additional RNase H-like domain sequences in Uniprot. JSON files were downloaded for the resultant data. After all the data was collected, the Whole Genome Shotgun (WGS) sequenced assemblies were removed from the InterPro datasets. After removing WGS sequences, the dataset was reduced to 1240 entries that included RNase H-like domains of THAP9-like proteins as well as some other proteins which bear similarity with the domain.

### Data organisation

The collected dataset (FASTA file) was arranged as per taxonomic hierarchy. The metadata for taxonomic details of each dataset was collected from NCBI Taxonomy Database (https://eutils.ncbi.nlm.nih.gov/entrez/eutils/efetch.fcgi?db=taxonomy&id={tax_id}&retmode=xml) (O’Leary et al. 2024). The dataset was arranged in the order – “Kingdom”, “Phylum”, “Class”, “Order”, “Family” and “Genus”. CD-HIT was performed on the dataset at 80% similarity to remove redundant entries (Fu et al. 2012). For this, the FASTA files of each dataset were disassembled for each individual species (using a custom python script). After performing CD-HIT for all the species individually, all the sequences were reassembled. This led to the reduction of the dataset to 722 sequences. After that all the non RNaseH-like domains were removed and the sequences got reduced to 409.

### Data visualisation

The MAFFT algorithm was used for creating Multiple Sequence Alignment (MSAs) with the FASTA files, to identify conserved regions as well as possible class-specific mutations (Katoh and Standley 2013). Jalview was used for visualization of the MSA, with a colour scheme based on the nature of amino acid as per the Clustal colours (Waterhouse et al. 2009).

**Clustal X Default Colouring**
| **Category** | **Colour** | **Residue at position** | **{ Threshold, Residue group }** |
| --- | --- | --- | --- |
| Hydrophobic | BLUE | A,C,I,L,M,F,W,V | {>60%, WLVIMAFCYHP} |
|  |  | C | {>60%, WLVIMAFCYHP} |
| Positive charge | RED | K,R | {>60%,KR},{>85%, K,R,Q} |
| Negative charge | MAGENTA | E | {>60%,KR},{>50%,QE},{>50%,ED},{>85%,E,Q,D} |
|  |  | D | {>60%,KR}, {>85%, D,E,N},<br>{>50%,ED} |
| Polar | GREEN | N | {>50%, N}, {>85%, N,D} |
|  |  | Q | {>60%,KR},{>50%,QE},{>85%,Q,T,K,R} |
|  |  | S,T | {>60%, WLVIAMFCYHP}, {>50%, TS},<br>{>85%,S,T} |
| Cysteines | PINK | C | {>85%, C} |
| Glycines | ORANGE | G | {>0%, G} |
| Prolines | YELLOW | P | {>0%, P} |
| Aromatic | CYAN | H,Y | {>60%, WLVIAMFCYHP}, {>85%,<br>W,Y,A,C,P,Q,F,H,I,L,M,V} |
| Unconserved | WHITE | any / gap | If none of the above criteria are met |

Using the MSA created by MAFFT, phylogenetic trees were created using IQ Tree which uses Maximum Likelihood (Nguyen et al. 2015). The parameters were kept as default and 1000 bootstraps were performed for both the datasets. The resultant tree was then visualized, coloured and analyzed using the itol online tool.

### Sequence clustering

After performing CD-HIT, clustering of data was done using DIAMOND at 40% similarity threshold to get representative sequences based on similarity (Buchfink et al. 2026). Therefore, two rounds of clustering were performed: first based on species using CD-HIT at 80% similarity, and the second clustering was done based only on sequence using DIAMOND at 40% similarity. The resultant dataset had 296 clusters.

### Structure based alignment

Multiple Structural Alignment (MSTA) was performed to help find distant homologs between the clusters that may not have been observable by just sequence-based alignments. First using the centroid of each cluster, AlphaFold structures of the domains were generated (Abramson et al. 2024). For clusters of each dataset, MSTA was performed using MUSTANG and Pymol (Konagurthu et al. 2006). MUSTANG and Pymol did not work for clustering the RNase H-like domains since the structures were too distant for them to align.

### Site directed Mutagenesis

Primers were designed for generating domain truncated mutants using NEB Base changer tool^TM^. Site directed mutagenesis, using Phusion polymerase (F-530S, Thermoscientific) and KLD enzyme (M0554S, NEB), was performed as per manufacturer’s instructions. All mutant constructs were verified by Sanger’s sequencing at CCAMP, India.

### *In-vivo* Integration assay

Cg4-Neo reporter plasmid (G418 resistance gene flanked by the TIRs of Cg4 P-element vector, transcribed by SV40 promoter) was co-transfected with hTHAP9 (wild type or mutants separately, cloned in pcDNA3.1(+)) or negative control (pBluescript empty vector) in HEK293 cells (0.4-0.5 × 10^6^ cells per well of a 6-well plate) at 70-90% confluency with Turbofect ^TM^ (R0531, Thermoscientific) transfection reagent, as per the instruction manual. Cg4-Neo, pBluescript vectors were generous gifts from Prof. Donald Rio, UC Berkeley.

Briefly, a total of 2 ug of plasmid DNA was transfected per well: 50 ng of CgNeo4, 1 ug hTHAP9 (wild type or mutants separately) /negative control and made up with pBluescript empty vector (Sharma et al. 2021). After 48 hours, the cells from each well were harvested and seeded to separate 10 cm dishes and allowed to adhere for 24 hours at 37 °C in a CO_2_ incubator. After 24 hours, media supplemented with G418 (0.5 mg/mL, TCI, Cat. no. G0349) was added to the cells. and selected for 2-3 weeks. The G418-resistant colonies were fixed using methanol, stained with crystal violet and counted. The relative integration activity was computed as a percentage of activity of each mutant to the wild type hTHAP9. The values are normalised with wild type hTHAP9.

Relative Integration Activity= (No. of G418 resistant colonies for mutants *100)/ (No. of G418 resistant colonies for wild type hTHAP9)

Significant values were determined using a one-way ANOVA method followed by Dunn Sidak test (Sharma et al. 2021).

## Results

### Homologs of the RNase H-like domain of hTHAP9 are widely distributed in animals from Cnidaria to Chordates

To investigate the evolutionary conservation of the RNase H-like domain of hTHAP9, homologous sequences were downloaded from UniProt (11466 entries, accessed in April 2026), trimmed for RNase H-like domain boundaries by using the corresponding boundary residues in hTHAP9 (186-679) and arranged taxonomically as per the hierarchy of Kingdom, Phylum, Class, Order, Family and Genus. CD-HIT was performed to remove redundant entries. This extensive analysis led to the identification of 722 entries which included sequences which are THAP9-like, *Drosophila* P-element-like or transposase-like as well as other proteins like p-32 hm, p-15 hm, p1 ap, etc. The latter proteins which are not related to THAP9/P-element like/transposases were removed so that the final dataset contained 409 entries which were subjected to Multiple Sequence Alignment using MAFFT.

We observe that homologs of the hTHAP9 RNase H-like domain are present in members of Animalia: from Cnidaria (Anthozoa, Hydrozoa), Platyhelminthes (Macrostomida), Arthopoda (Insecta, Arachnida, Copepoda, Malacostraca), Echinodermata (Asteroidea), to Cephalochordata (Leptocardii) and Chordata (Hyperoartia, Myxini, Actinopteri, Amphibia, Lepidosauria, Reptilia, Aves, Mammalia). Fig 4 shows the overall percentage distribution of organisms possessing this domain with highest representation in Insecta and Arachnida (Fig. 4a) and low representation in Hyperoartia, Leptocardii and Macrostomida (Fig. 4b).

**Fig 4:**
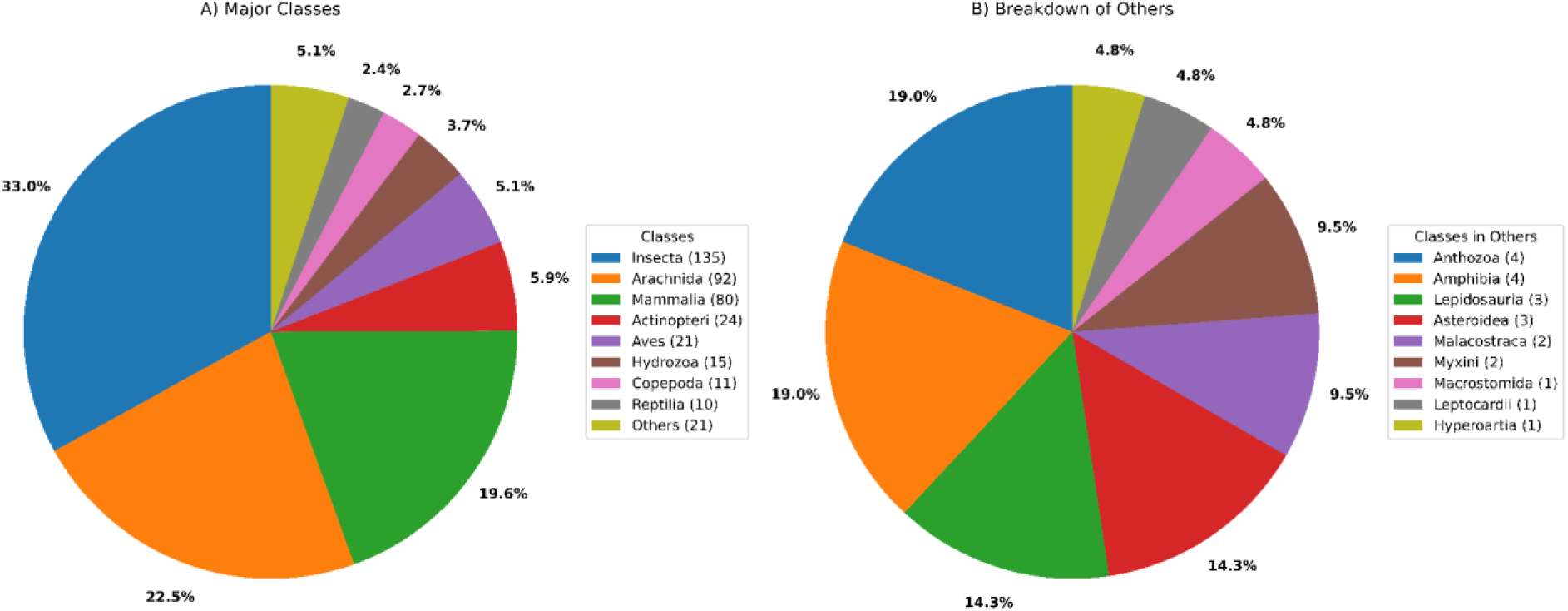
Homologs of RNase H-like domain of hTHAP9 are spread across invertebrates and vertebrates with (a) maximum representation in insects (33%) within invertebrates and mammals (∼20%) within vertebrates (b) Lower occurrence is found in platyhelminthes (Macrostomida), cephalochordates (Leptocardii) and jawless fishes (Hyperoartia)

### Length distribution of the RNase H-like domain of hTHAP9 homolog

The lengths of the homologs of the hTHAP9 RNase H-like domain were then investigated (Fig 5).The largest RNase H-like domain is 507 amino acid residues, found in Common Lancelet, belonging to Leptocardii. The average length of the RNase H-like domain is longer in groups having higher representation i.e. the vertebrates (Reptilia, Aves, Mammalia) than the invertebrates (Insecta and Arachnida) (Fig 5).Thus, the average length of the domain increases in a class-specific manner, indicating hierarchical increase domain size during evolution.

**Fig 5:**
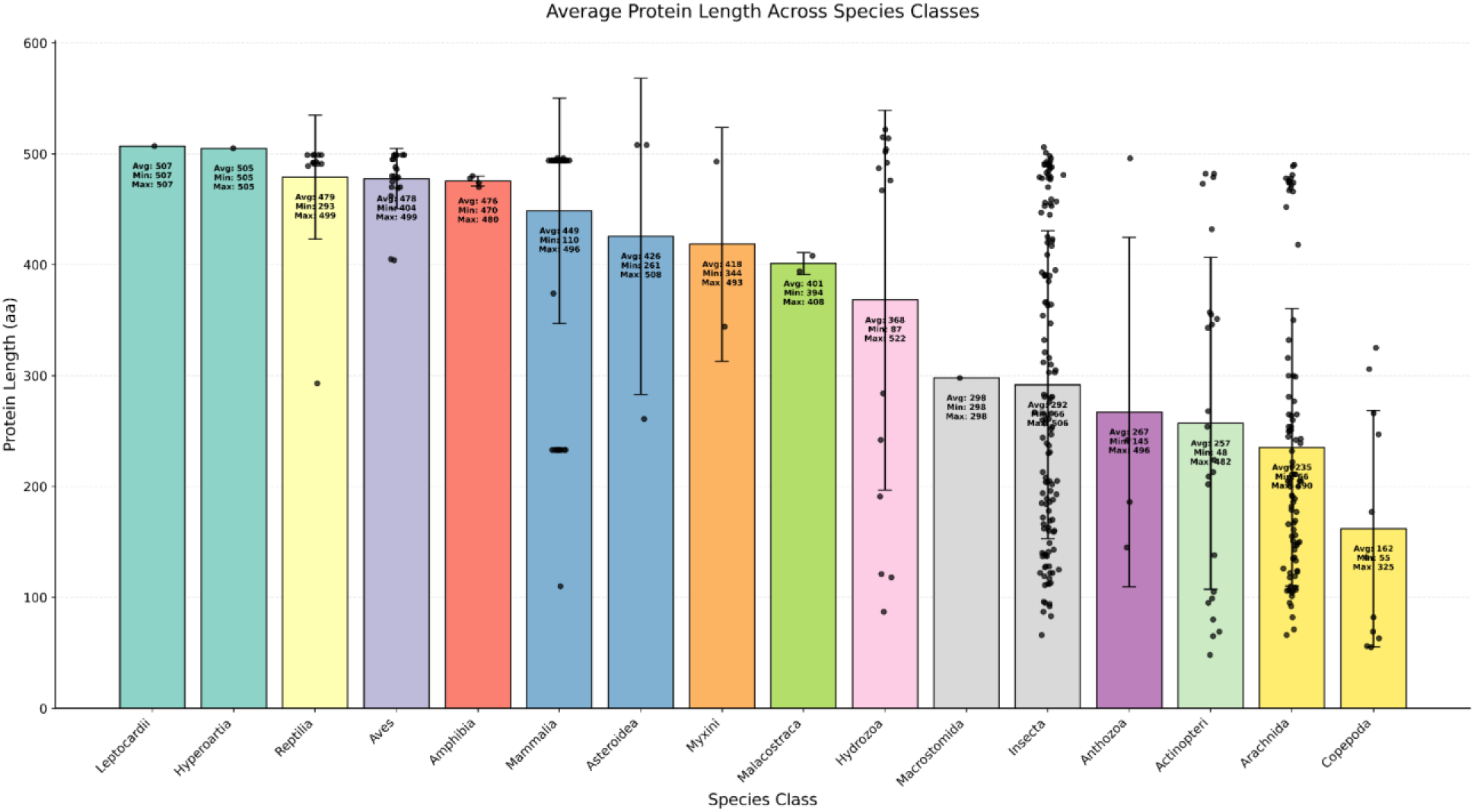
The average length of the RNaseH-like domain increases from Arthropoda to Vertebrates. Individual data points corresponding to lengths of individual domains are overlaid on the bars with the error bars representing standard deviation, X-axis representing the organism class and Y-axis representing the average length of the RNaseH-like domain.

### Phylogenetic analysis of RNase H-like domains in transposases similar to THAP9 follows pattern of evolution

Multiple sequence alignment was initially performed with all homologs of THAP9 RNase H-like domains which included both transposase proteins as well as non-transposase proteins. It was observed that the non-transposases shared only certain hydrophobic regions (Suppl. Fig 3) which are similar but not identical; hence they were removed from the list. The non-transposase proteins included uncharacterised proteins or those with unknown function as well as HMCN1 (involved in cell-cell adhesion), NADH dehydrogenase [ubiquinone] 1 alpha subcomplex subunit 12 (accessory subunit of mitochondrial membrane respiratory chain), 2-oxo-4-hydroxy-4-carboxy-5-ureidoimidazoline decarboxylase (facilitates the production of allantoin),etc. whose functions are not related to RNase H-like activity and hence removed. MSA was then performed with transposase derived proteins.

To study the evolutionary history of the homologs of RNase H-like domains, phylogenetic trees were constructed using MSAs (Multiple Sequence Alignment) created by MAFFT after performing CD-HIT. It was observed that the pattern of occurrence of this domain followed the order of evolution: Cnidaria, Arthropoda, Mollusca, Brachiopoda, Chordata. Organisms from Reptilia, Aves, Chordata clustered together thus illustrating higher conservation among member organisms of the same class (Fig. 8). However, few organisms from class Arthropoda interspersed within with Pisces and Amphibia.

### Clustering of RNase H-like domain of transposases similar to THAP9

The homologs of the THAP9 RNase H-like domain were then clustered (using DIAMOND, clustering was performed at 40% sequence similarity) to club organisms possessing similar domains in one group. In a cluster, the centroid is the representative sequence or structure that most accurately captures the characteristics of every other member. This representative sequence was extracted for each group, thereby reducing redundancy and the volume of the datasets. This led to the identification of 296 distinct clusters from the 409 sequences (Suppl. Table 1). The maximum number of sequences were clustered into a single entry representing mammals (centroid as Philippine tarsier, 66 members). This suggested that there was strong class-specific sequence conservation amongst these mammalian members.

The structures of the representative sequences obtained after DIAMOND clustering were then predicted using AlphaFold. The majority of the representatives (Suppl. Fig. 1) were observed to possess the characteristic RNaseH like fold: five β sheets, often sandwiched by α helices which also corresponded to the high confidence regions (marked in shades of blue) in the AlphaFold predicted structures (Majorek et al. 2014).

Phylogenetic analysis (Suppl. Fig 10) of the centroid sequences illustrate that the members of Arthropoda do not cluster together but instead form multiple branches in the phylogenetic tree. The arthropods are a diverse set of organisms with varied classes, which have diverged more deeply than mammals (Pisani et al. 2004; Lee and Beck 2015) .

To investigate if there was structural conservation amongst the individual cluster members, Multiple Structural Alignment (MSTA) was performed using MUSTANG. However, the homologs of THAP9 RNase H-like domain could not be aligned to a common core. The possible reasons could be the variability in the insertion domain in individual RNase H-like domains or the presence of incomplete domains which lacked all secondary structure elements.

### Multiple sequence alignment of RNase H-like domains illustrate conserved amino acid residues across all classes

Multiple sequence alignment was performed with the RNase H-like domains of transposase derived proteins. There are several proteins which have an incomplete RNase H-like domain (Fig 6) wherein some may have only the N-terminal end of the domain [e.g. Green monkey (A0A0D9QWP2)], some only the C-terminal end [e.g. Tuatara (A0A8D0H3H5), Orbiculate cardinalfish (A0A672YJQ0)] while there are some mammals which do not have the both N-and C-terminal ends but only possess the insertion domain (ID) [e.g.Long-eared bat (Q59AC6), European hare (Q59AC5), European polecat (Q59AC1), dog (Q59AC2), cat (Q59AC3), Western gorilla (Q59AB1), Bornean orangutan (Q59AB4), Lar gibbon (Q59AB5), Common Squirrel monkey (Q59AB7), Ring-tailed lemur (Q5AB8)].

**Fig 6:**
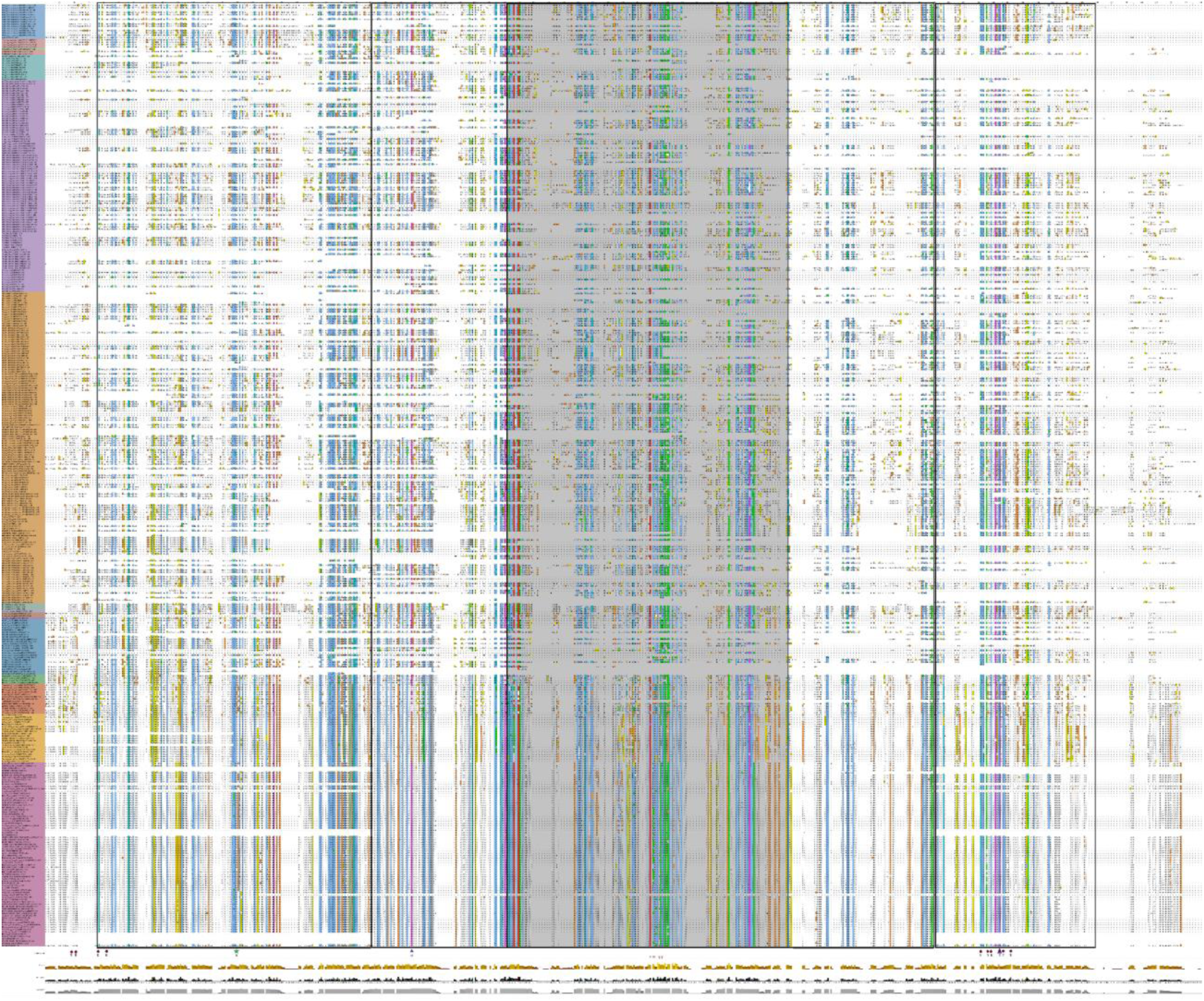
MSA of proteins with RNase H-like domains from transposable element derived proteins with YREK and YKEFR motifs (brown circles), K282 (green star) and catalytic residues D and E (blue triangle). The MSA can be divided into 3 sections with the first and last section harbouring residues which are not identical but similar (blue hydrophobic patches). The intervening middle region corresponds to the ID in hTHAP9 (region marked in grey).

It is interesting to note that the IDs within the RNase H-like domain are class specific i.e., they are similar within members of the same class but differ when compared to other classes. Notably, some homologs in classes like Actinopteri consist of the N and C-terminal parts of the RNaseH-like domain, but do not have the intervening ID while there are some homologs which lack both the ID as well as the C-terminal part of the RNase H-like domain (Fig 7).

**Fig 7:**
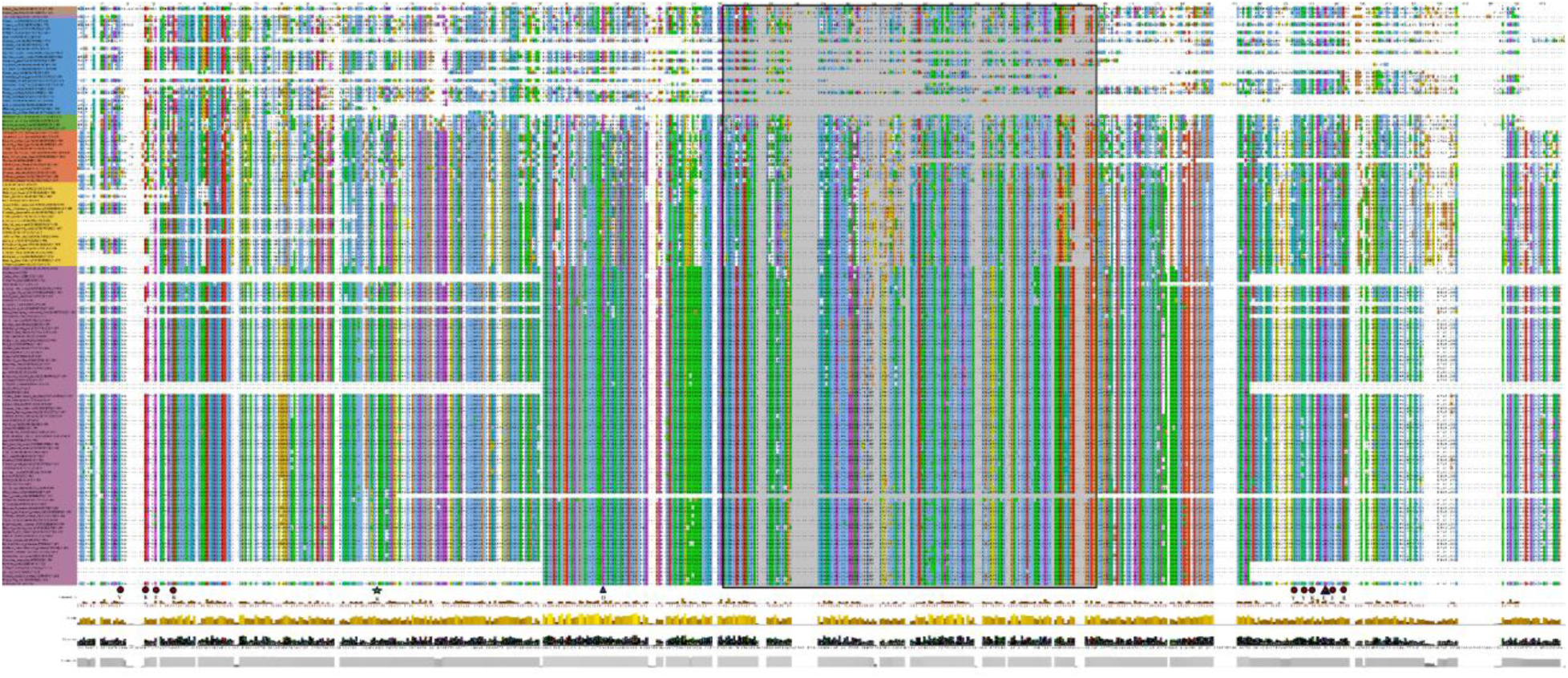
MSA of RNase H-like domains from transposable element derived proteins of vertebrates; Actinopteri - blue, Amphibia-green, Aves-yellow, Hyperortia-lavender, Mammalia-mauve, Myxini-brown, Reptilia-terracotta with with YREK and YKEFR motifs (brown circles), K282 (green star) and catalytic residues D and E (blue triangle). The intervening middle region corresponds to the ID in hTHAP9 (region marked in grey).

**Fig 8:**
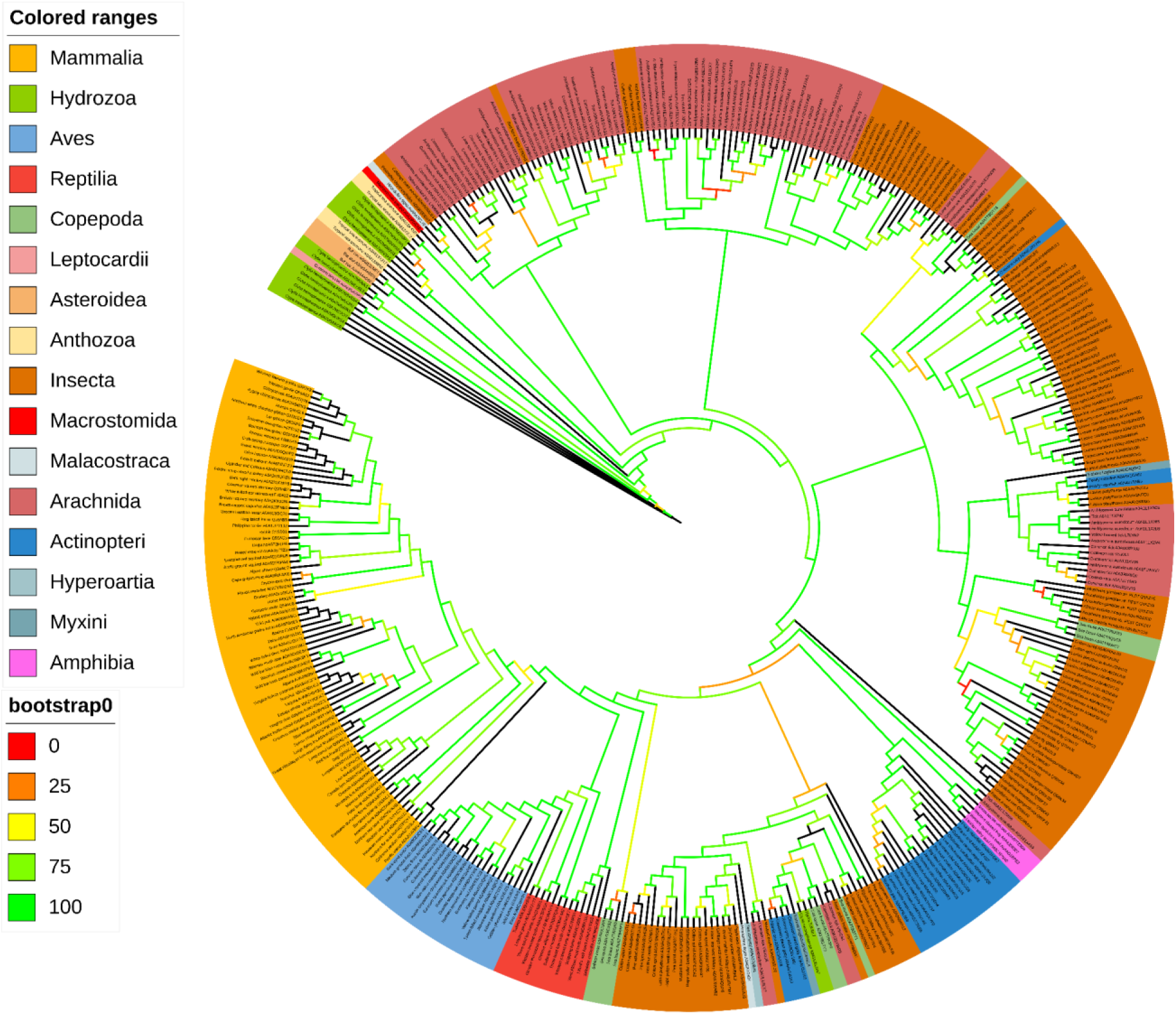
Phylogenetic analysis of homologs of THAP9 RNAse H-like domain protein sequences after CD-HIT: The sequences are aligned using MAFFT and the tree is generated using IQ TREE showing the presence of the domain from Hydrozoa (Cnidaria) to Mammalia (Chordata).

### Phylogenetic analysis of YREK and YKEFR residues across vertebrate species along with K282

The superposition of RNase H-like domains of DmTNP and hTHAP9 (Rashmi et al. 2024) illustrated that the two of the catalytic residues (D303 and E531) within the DDE catalytic triad of DmTNP aligned with D374 and E613 of hTHAP9. Interestingly, it has previously been shown that D374 and E613 of hTHAP9, are important for catalysizing the DNA excision and integration (Sharma et al. 2021). However, the third DmTNP catalytic residue (D230) aligns with K282 of hTHAP9 (fig 2). Interestingly, this Lysine residue was conserved across all vertebrate THAP9 homologs. Thus, we decided to explore the role of this conserved Lys in mobilisation of *Drosophila* P-element TIRs.

Further, the YREK (Y200, R203, E207, K213) and YKEFR (Y601, Y605, K606, E613, F615, R620) motifs present on either end of the RNase-H-like catalytic domain of hTHAP9, showed class-wise evolutionary conservation (Fig. 9). Interestingly, both these motifs are absent in RNaseH-like domain of DmTNP suggesting that they may have a specific role in regulation of catalytic activity in hTHAP9 and its homologs. Notably, the YKEFR motif encompasses the catalytic glutamate (E613) residue. With the exception of K213 and F615, all the residues show class specific divergence (Table 1 and Fig 9). Certain homologs (in several members of Actinopteri and Mammalia) lack either the YKEFR motif or both the YREK and YKEFR motifs while some homologs (from Aves) have an incomplete YREK motif where only E and K are retained. Moreover, very few (5) homologs lack the YREK motif but retain the YKEFR motif. Thus, we also decided to investigate if these motifs played a role in DNA integration by hTHAP9.

**Fig 9:**
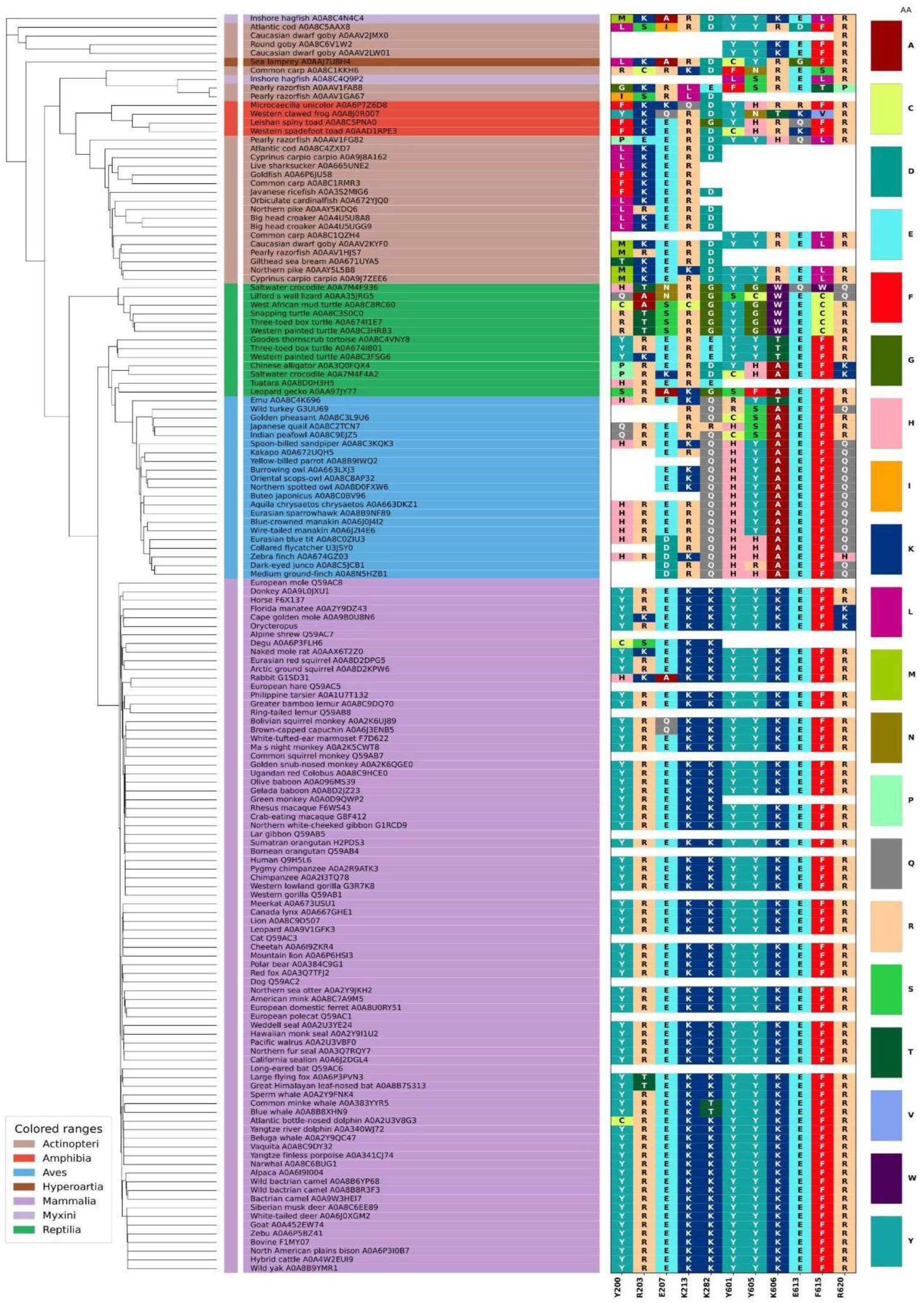
Phylogenetic analysis of RNase H-like domains from transposable element derived proteins of vertebrates. YREK (Y200. R203, E207, K213) and YKEFR (Y601, Y605, K606, E613, F615, R620) motifs along with K282 illustrates their class-specific conservation

**Fig 10:**
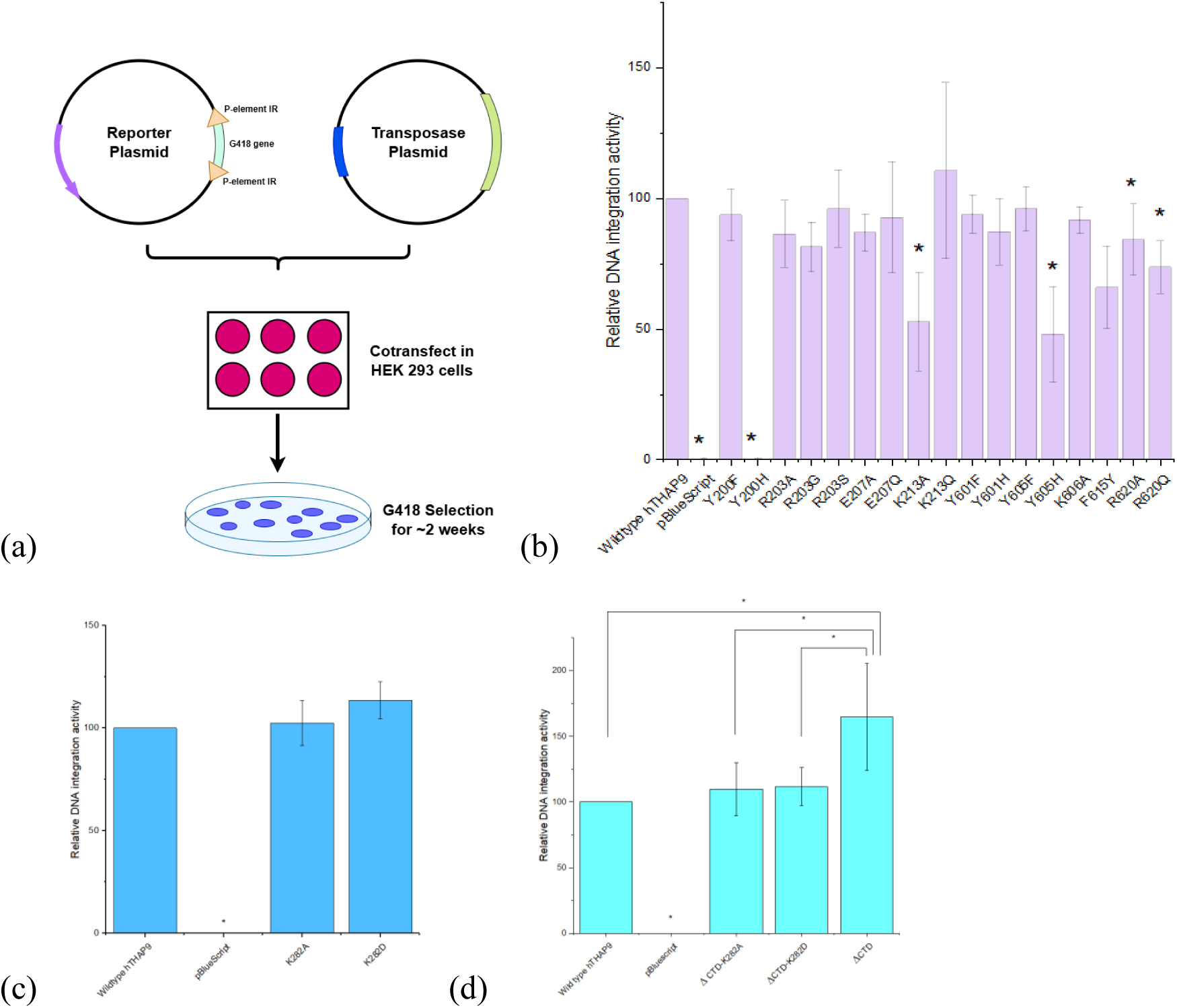
(a) Schematic of the DNA integration assay. Relative DNA integration activity of the (b) hTHAP9 point mutants from YREK and YKEFR motifs (c)hTHAP9 K282 mutants (d) hTHAP9 double mutant of K282A or K282D with ΔCTD.

**Table 1:** List of amino acid substitutions in YREK, YKEFR motifs, K282.

| **Motif residues** | **Amino acid substitutions** | **Substitution based on physicochemical properties** | **Evolutionarily observed substitution** |
| --- | --- | --- | --- |
| Y200 | Y200F, Y200H | Y200F | Y200H (Aves) |
| R203 | R203A, R203G, R203S | R203G, R203A | R203A (Reptilia), R203S (Actinopteri) |
| E207 | E207A, E207Q | E207A | E207A (specific members of Actinopteri, Reptilia and Mammalia), E207Q (specific members of Actinopteri) |
| K213 | K213A, K213Q | K213A, K213Q |  |
| K282 | K282A, K282D | K282A | K282D (Actinopteri, Amphibia and Reptilia) |
| Y601 | Y601F, Y601H | Y601F | Y601H (Actinopteri, Amphibia, Reptilia and Aves) |
| Y605 | Y605F, Y605H | Y605F | Y605H (Aves) |
| K606 | K606A | K606A | K606A (Reptilia and Aves) |
| F615 | F615Y | F615Y |  |
| R620 | R620A, R620Q | R620A | R620Q (Aves) |

### Mutational analysis of conserved motifs in the RNaseH-like domain of hTHAP9

To investigate the roles of the conserved YREK and YKEFR motifs as well as the Lys282 residue in the catalytic activity of hTHAP9, individual point mutants were created using site-directed mutagenesis on the wildtype template. Table 1 contains the list of targeted residues along with the substituted amino acid.

Some of the amino acid substitutions were designed based on the physicochemical differences (Table, e.g., charge, hydrophobicity, aromaticity, side-chain size etc) between the native and substituted residues. For example, substitution of tyrosine to phenylalanine (Y→F) retains the aromatic ring and hydrophobic character of tyrosine while eliminating the hydroxyl group, thereby allowing the contribution of the hydroxyl group to be assessed. Similarly, substitution of lysine or arginine with alanine (K/R→A) removes the positively charged side chain and substantially reduces side-chain size, allowing the importance of positive charge and side-chain bulk to be evaluated. By introducing such changes in properties, the role of specific physicochemical properties of individual residues for the structural and functioning of hTHAP9 can be investigated.

Substitutions were also made based on observations made in the phylogenetic analysis (MSA in Fig.9, Table 1) wherein certain THAP9 residues appeared to be conserved in particular taxonomic lineages and underwent class-specific substitutions across different vertebrate classes (Fig, 9). Thus the substitutions were selected to reproduce amino acid states that have naturally occurred and been retained during the course of THAP9 evolution. Such substitutions allow the functional consequences of naturally occurring evolutionary variation to be examined in the context of the hTHAP9 protein. For example, as per the MSA, Y200 (in hTHAP9) aligned with histidine in all Aves; therefore, the Y200H mutation was generated in hTHAP9 to reproduce this evolutionarily observed substitution. Similarly, R203(in hTHAP9) aligned with Serine in Actinopteri, resulting in the selection of the R203S mutation; the Y601H mutant was made because Y601(in hTHAP9) aligned with His in Actinopteri, Amphibia, Reptilia and Aves while the R620Q mutant was made because R620(in hTHAP9) aligned with glutamine in Aves, and the mutation.

K282 (in hTHAP9) aligned with Aspartate in Actinopteri, Amphibia and Reptilia; therefore, K282D was selected to investigate the effect of introducing this naturally occurring variant state into hTHAP9. Interestingly, given that K282 (in hTHAP9) superimposes with an Asp in the catalytic triad of DmTNP (Fig. 2), the K282D mutant also represented a mutant hTHAP9 with a modified DmTNP-like catalytic triad. Thus, these substitutions were selected directly from residue variation observed in the MSA and were used to determine whether amino acid states naturally present in other vertebrate lineages can have any functional relevance in hTHAP9.

### Characterization of conserved motifs in the RNaseH-like domain of hTHAP9

We next investigated whether the YREK and YKEFR motifs as well as K282 were required for DNA integration by performing cell-based integration assays with all the generated point mutants (Table 1). The ability of the transposase source (wild type or mutant) to excise and then integrate the G418 resistance cassette flanked by P-element TIRs was assessed by screening and staining G418-resistant colonies (described in the Methods section). The relative DNA integration activity (Suppl. Table 2, Sharma et al. 2021) was then calculated. Overall, the relative integration activities of the mutants were 70-100% compared to wildtype hTHAP9 (Suppl. Table 2).

*YREK motif*. The residues of the YREK motif appeared to have varying importance in catalysis. While DNA integration activity is unaffected by the substitution of Y200 with F, changing the residue to H completely abolishes the activity. Histidine, harbouring a five-membered imidazole ring, has a smaller ring size than Tyr, which can affect its surroundings and interfere with its bonding partners (Lanzarotti et al. 2011). Substitutions of R203 (to A, G, S) and E207 (to A, Q) do not have much effect, indicating that these residues probably do not contribute much to DNA integration.

On the other hand, K213 appears to be an important residue since changing it to A significantly lowers its DNA integration ability (∼50% of wildtype), whereas changing it to Q increases its DNA integration ability slightly (though not significantly). It is tempting to speculate that the observed importance may be because Lysine is a positively charged amino acid which can directly interact with the negatively charged DNA.

*YKEFR motif*. The residues of the YKEFR motif also appeared to have varying importance in catalysis. Both Y600 and K606 are tolerant to mutations and are probably not essential for DNA integration. However, Y605 like Y200 appears to be important for DNA integration. Interestingly, much like Y200, DNA integration activity is unaffected by the substitution of Y605 with F, while changing the residue to H decreases the activity(∼50% of wildtype activity).

E613 has previously been shown to be an important catalytic residue (Sharma et al. 2021) as mutation to Gln severely reduced DNA integration (7.5% of wildtype activity). Substitutions in both F615 and R620 lead to moderate decrease in integration activity (∼60-80% of wildtype activity, Suppl. Table 2), suggesting that these residues contribute to catalytic activity but are not indispensable.

*Lys 282*: Surprisingly, unlike our hypothesis, K282 does not appear to have a prominent catalytic role since its substitution does not significantly enhance hTHAP9 activity as compared to wild type. This suggests that even though the residue overlaps with a catalytic Asp in the DmTNP structure, it may not contribute much to catalysis in hTHAP9. Thus, it is possible that hTHAP9 may not have a canonical DDE catalytic triad like DmTNP. Alternatively, other unexplored acidic residues close to K282 may prove to be the elusive third catalytic residue.

Lys282 was also mutated on the background of the hyperactive CTD truncation mutant of hTHAP9 (studied in Chapter 2; showed enhanced DNA integration activity): these double mutants also had activity similar to wild type suggesting that K282 somehow inhibited the hyperactivity associated with CTD truncation. Thus, it is tempting to speculate that although K282 appears to be dispensable for DNA integration in the full length hTHAP9, it may influence the conformational or regulatory state of hTHAP9 that results in hyperactive phenotype following CTD removal.

Overall, these mutational analyses identify **Y200, K213** in the YREK motif and **Y605, E613, F615, R620** in the YKEFR motif as residues that make important contributions to hTHAP9-mediated integration, whereas **R203, E207** and **Y601, K606** appear to play comparatively minor roles under the conditions tested.

## Discussion

The RNase-H fold is considered to be one of the most ancient protein folds which originated from viruses but has been adapted by diverse proteins in all kingdoms of life. These include transposases, Piwi nucleases, RecQ, RAG1 etc. which are responsible for diverse processes like transposition, RNA silencing, helicases, recombination respectively (Moelling et al. 2017).

The inclusion of proteins in the dataset such as HMCN1, NADH dehydrogenase subunits and metabolic enzymes showcases one of the limitations of computational domain prediction. RNase H-like fold is exhibited by numerous unrelated protein families that perform completely different biological functions. However sequence-based domain searches often retrieve proteins sharing only sequence similarities rather than common evolutionary ancestry. This observation reinforces the need to combine sequence similarity, functional annotation and phylogenetic context while studying ancient protein folds (Majorek et al. 2014). Similar conclusions have been reported for RNase H-like nucleases, where proteins with similar tertiary structures perform remarkably diverse biological roles (Hyjek et al. 2019).

An unexpected observation in this study is the appearance of proteins possessing only partial RNase H-like domains. Several vertebrate proteins retain only the N-terminal end, others preserve only the C-terminal end, whereas some mammals contain predominantly the insertion domain (ID).Several explanations may account for these observations.Firstly, there can be incomplete genome annotations or fragmented protein predictions, particularly among non-model organisms. Secondly, these proteins could represent partially active or inactive transposases, losing their catalytic activity after recruitment into host cellular pathways as observed in the case of SETMAR and RAG1, where residues in the catalytic domain have been modified without causing much loss to its structural architecture (Shaheen et al. 2010; Kapitonov and Jurka 2005).Third, relaxation of selective pressure due to loss of transposition ability, may have led to progressive loss of regions which are no longer required for DNA transposition while retaining other domains necessary for protein–protein or protein–DNA interactions. Moreover, proteins which retain only the IDs may have acquired independent functional significance, not being limited only to catalysis. Further experimental validation will be required to distinguish between annotation artefacts and genuine evolutionary intermediates.

Previous studies have established D374 and E613 as catalytic residues essential for hTHAP9-mediated DNA excision and integration (Sharma et al. 2021). Structural comparison with DmTNP transposase also indicated that these hTHAP9 residues overlapped with D303 and E531 in the catalytic triad of DmTNP. However, the third catalytic residue (D230) of DmTNP appeared to overlap with K282 in hTHAP9 (Rashmi et al. 2024). Although we had speculated that substituting the K282 residue with an Asp (or Ala) may enhance hTHAP9’s ability to integrate DNA, our results suggest otherwise. The absence of significant increase in integration activity suggests that structural equivalence does not necessarily imply functional equivalence. This finding also suggests that hTHAP9 may have evolved a catalytic mechanism distinctly different from that of its *Drosophila* counterpart, with K282 likely serving a non-catalytic role. Domesticated transposases often exhibit modifications in their catalytic machinery while acquiring some host-specific cellular functions. The absence of a conventional DDE catalytic triad in hTHAP9, despite the retention of measurable DNA integration activity, suggests that the protein might have undergone evolutionary adaptation rather than complete loss of catalytic function. This may represent one of the several molecular signatures associated with the domestication of hTHAP9. Alternatively, other acidic residues (including the previously identified important catalytic residue D304; Sharma et al. 2021) close to K282 may prove to be the elusive third catalytic residue. Additional structural and biochemical studies are required to illustrate how catalysis is achieved in the absence of a canonical catalytic triad.

This study also analyzed the evolution of conserved residues on either termini of the catalytic RNase H-like domain. Most residues of the YREK and YKEFR motifs display class-specific divergence, suggesting that these regions have evolved under lineage-specific selective pressures. Among the other residues, K213 and F615 remain slightly more conserved than others, implying stronger functional constraints. Another interesting observation is that organisms lacking YKEFR still retain YREK, but no species lacking YREK preserves YKEFR in the dataset analysed. This evolutionary pattern suggests that YREK may constitute an ancestral structural element while YKEFR evolved later during vertebrate diversification. Future functional studies are required for confirming these hypotheses. The conservation of YREK and YKEFR as intact sequence motifs across mammalian THAP9 homologs, in contrast to their absence in other vertebrate classes, suggests that these sequences may represent mammal-specific evolutionary features. Their strong conservation within mammals indicates that they may have acquired structural or functional significance during mammalian evolution.

The dramatic reduction in integration following mutation of Y200H, while Y200F remains fully active, reinforces the necessity of maintaining an aromatic residue more than retention of the hydroxyl group. Though Histidine is aromatic, it is highly polar and hydrophilic, thereby participating in other interactions. Aromatic residues frequently stabilize protein cores through π-stacking interactions when placed at specific orientation, or contribute to ligand recognition via base stacking (Burley and Petsko 1985; Hillier et al. 1998). Similar aromatic requirements have been reported in Hermes transposase, where substitution of conserved aromatic residues disrupts DNA binding despite having catalytic residues intact (Alison B. Hickman et al. 2014).

K213 is perhaps the most intriguing residue identified in this study, whose mutation to specific amino acids can significantly reduce or increase integration modestly. Lysine-to-glutamine substitutions are employed as mimics of lysine acetylation because glutamine neutralizes the positive charge while preserving side-chain geometry (Wang and Hayes 2008). Although speculative, the increased activity observed for K213Q raises the possibility that neutralization of this residue weakens electrostatic interactions with DNA, facilitating product release after strand transfer and thereby improving catalytic turnover. Similar regulatory effects of lysine acetylation have been reported for several DNA-binding proteins (Choudhary et al. 2014).

Mutation of Y605 mirrors the behaviour of Y200, suggesting that aromatic interactions contribute to optimal DNA integration. Replacement of F615 with tyrosine moderately reduces activity, indicating that maintenance of a hydrophobic aromatic environment is favourable at this position. Similarly, R620 contributes to efficient DNA integration without being absolutely dispensable. Altogether these observations demonstrate that residues apart from the canonical DDE catalytic triad significantly influence DNA integration efficiency, most likely by stabilizing substrate binding, positioning catalytic residues or maintaining active-site architecture.

## Conclusions

This study demonstrates that the RNase H-like domain of hTHAP9 and its homologs has undergone substantial evolutionary diversification while retaining its fundamental catalytic architecture. Expansion of class-specific insertion domains, divergence of non-catalytic motifs and selective conservation of structurally important residues, among others, illustrate how transposase derived genes evolve to possess novel biological functions without disrupting their ancestral RNase H-like scaffold. Integration of evolutionary analyses with functional mutagenesis identifies Y200, K213, Y605, F615 and R620 as previously unrecognized contributors to hTHAP9 activity and provides new insights into its molecular evolution. These findings widens our understanding of how ancient transposases could have been rewired during vertebrate evolution and establish a framework for future structural and biochemical investigations into hTHAP9 function.

